# FLiPPR: A Processor for Limited Proteolysis (LiP) Mass Spectrometry Datasets Built on FragPipe

**DOI:** 10.1101/2023.12.04.569947

**Authors:** Edgar Manriquez-Sandoval, Joy Brewer, Gabriela Lule, Samanta Lopez, Stephen D. Fried

## Abstract

Here, we present FLiPPR, or FragPipe LiP (limited proteolysis) Processor, a tool that facilitates the analysis of data from limited proteolysis mass spectrometry (LiP-MS) experiments following primary search and quantification in FragPipe. LiP-MS has emerged as a method that can provide proteome-wide information on protein structure and has been applied to a range of biological and biophysical questions. Although LiP-MS can be carried out with standard laboratory reagents and mass spectrometers, analyzing the data can be slow and poses unique challenges compared to typical quantitative proteomics workflows. To address this, we leverage the fast, sensitive, and accurate search and label-free quantification algorithms in FragPipe and then process its output in FLiPPR. FLiPPR formalizes a specific data imputation heuristic that carefully uses missing data in LiP-MS experiments to report on the most significant structural changes. Moreover, FLiPPR introduces a new data merging scheme (from ions to cut-sites) and a protein-centric multiple hypothesis correction scheme, collectively enabling processed LiP-MS datasets to be more robust and less redundant. These improvements substantially strengthen statistical trends when previously published data are reanalyzed with the FragPipe/FLiPPR workflow. As a final feature, FLiPPR facilitates the collection of structural metadata to identify correlations between experiments and structural features. We hope that FLiPPR will lower the barrier for more users to adopt LiP-MS, standardize statistical procedures for LiP-MS data analysis, and systematize output to facilitate eventual larger-scale integration of LiP-MS data.

## INTRODUCTION

Structural proteomics is an expanding subfield within the space of proteomics that aims to explore protein structure, dynamics, and stability in a global, unbiased manner. The field is defined by several emerging methods that convert structural information into mass, enabling mass spectrometry data to measure structural information on many distinct proteins in a complex mixture. Four leading structural proteomic methods include hydrogen-deuterium exchange (HDX),^1–3^ methionine oxidation methods (e.g., SPROX),^4,5^ fast photochemical oxidation of proteins (FPOP), ^6,7^ and limited proteolysis mass spectrometry (LiP-MS).^8–10^ In these methods, regions within proteins that are solvent-accessible are labeled, respectively, by deuterium at backbone amides, by oxidation at methionine from H2O2, by oxidation at any amino acid from HO radicals, or by cleavage from a sequence-permissive. ^11^ Crosslinking mass spectrometry (XL-MS) is another method that can provide rich structural information in the form of residue-residue contacts (which can serve as distance restraints),^12–15^ though it differs from the other methods in that it requires sequencing low-abundance crosslinked peptides from vast search spaces, creating a unique set of technical challenge.^16,17^

Amongst the labeling-based methods, LiP-MS has emerged as a popular structural approach because it has a few key advantages (e.g., residue-level resolution, proteome-wide coverage) without some drawbacks that affect other methods (e.g., need for specialized purpose-built instruments, need to identify rare low-abundance species, significant amplification of the search space). In its modern form (devised by Feng et al.), the experimentalist subjects a complex mixture of proteins to a pulse of proteolysis with a non-specific protease (typically proteinase K) under native conditions, causing solvent-accessible or unstructured portions within proteins to get cleaved (Figure 1A).^9^ Afterward, the sample is subjected to in-solution trypsin digest under denaturing conditions, which produces a mixture of tryptic and half-tryptic peptides that are sequenced by LC-MS/MS. The non-tryptic cut-site of each half-tryptic peptide reveals a residue that was solvent accessible in the parental protein. LiP-MS is a very accessible structural proteomic method because, in most samples, half-tryptic peptides are numerous, abundant, and don’t require specialized search settings to identify. Data-dependent acquisition (DDA)^8,18^ and data-independent acquisition (DIA)^10,19^ workflows have both been implemented. So far, it has been applied to probe a range of biological and biophysical questions at the proteome scale, such as metabolic rewiring in response to nutrients,^19^ aging in yeast,^20^ aging in rodents,^21^ thermostability,^22^ protein folding,^18,23^ among others.

**Figure 1.**
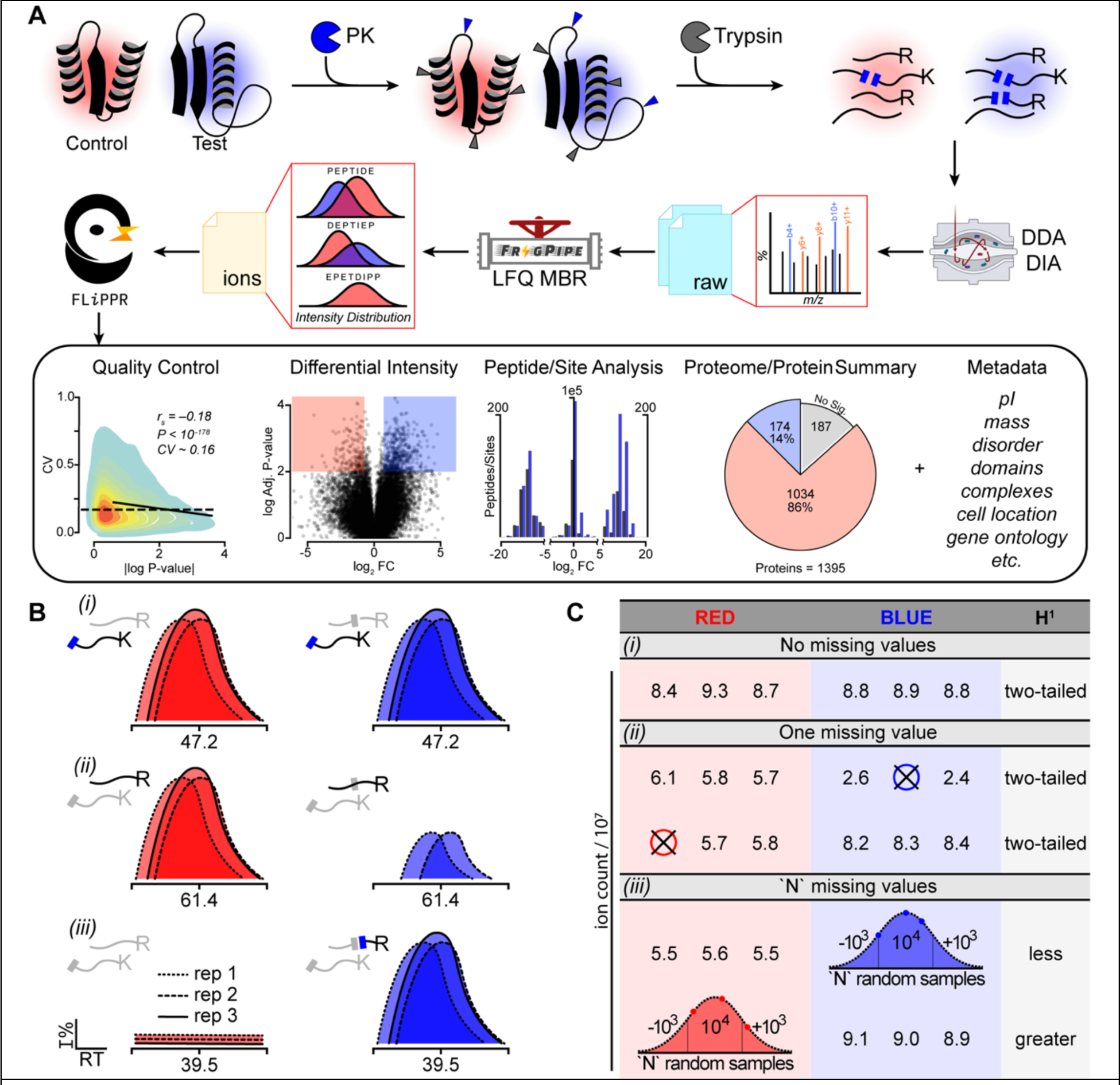
Summary of LiP-MS Workflow and Processing in FLiPPR. (A) The top row is a simple schematic of LiP-MS sample preparation, featuring a control condition (red) and a test condition (blue) simultaneously subjected to limited proteolysis (LiP) by proteinase K (PK) and then complete trypsinolysis. The second row represents data handling in the FLiPPR pipeline: raw mass spectra are searched and quantified in FragPipe, and the ions file is processed by FLiPPR, which produces a range of outputs featured in the third row. (B, C) How FLiPPR treats missing data at the ion level. Case *i*: If quantifications are available for an ion in all replicates of test and control, then averages are calculated, and the P-value is assessed with a two-tailed t-test. Case *ii*: If one quantification is missing in either the test or the control, the missing value is dropped, and the P-value is assessed with a two-tailed t-test comparing *n* to *n*–1 values. Case *iii:* If all the values are missing in either the control or the test (in the example shown in B, it is the control) and all the values are present in the other condition, then the ion is considered “all-or-nothing” (AON). The *n* missing values are filled by Gaussian imputation, then averages are calculated as in case i; P-values are calculated with a one-tailed test. All other permutations of missing data result in an ion being disqualified.

A LiP-MS study is usually designed as a quantitative experiment in which two (or more) closely related samples are generated and subjected to the same workflow; half-tryptic peptides that are present with significantly different abundances between two sample types represent locations associated with a structural change (strictly speaking, a change in proteolytic susceptibility) between the respective conditions. Most studies to date have employed area-under-the-curve label-free quantification (LFQ) to assess these abundance differences, with a few studies applying SILAC quantification or isobaric mass tag methods instead^5,8,24,25^ Some key advantages of LFQ-based quantification are its high dynamic range and remarkable ability to independently assess missing features in distinct conditions (or replicates), which has outsized importance in LiP-MS and can provide very insightful information if used judiciously.

The experimental details to prepare samples for LiP-MS have been carefully developed and advanced by Picotti and co-workers,^10^ which we have used with minor modifications in our adaptation of the method to study protein refolding (see Experimental Section in Supplementary Information). On the data analysis side, however, we have found that several developments from our lab have been valuable. While the relevance of these improvements was first realized in the context of our ongoing studies applying LiP-MS to protein folding, we believe they would be useful for LiP-MS studies in general, and here report a computational tool called FLiPPR (FragPipe LiP-MS Processor) to formalize our data analysis workflow and facilitate its adoption by the structural proteomics community.

First and foremost, FLiPPR implements a unique treatment of missing data tailored to LiP-MS. Sometimes, when a protein misfolds, core regions that were completely inaccessible to proteinase K can become proteolytically susceptible (as shown in the case of Figure 1A); this results in a situation in which a half-tryptic peptide will be detected only in samples containing the misfolded protein but will be absent in samples containing the native protein, resulting in missing data. A process is necessary to distinguish this scenario (where the missing data are informative (e.g., Figure 1B, case iii)) from scenarios where missing data should instead prevent a feature from being included in an analysis.

Secondly, LiP-MS is fundamentally a “peptide-centric” technique, and so data compression, propagation of error, and data disagreement must be considered from the level of ions to modified peptides to peptides, and – we argue here – ultimately to cut-sites (cf. Figure 2). Since most quantitative proteomics studies focus on the protein level and have an additional “protective” layer to buffer against noise by averaging across peptides, data processing for LiP-MS raises unique concerns not addressed by commercial software packages.

**Figure 2.**
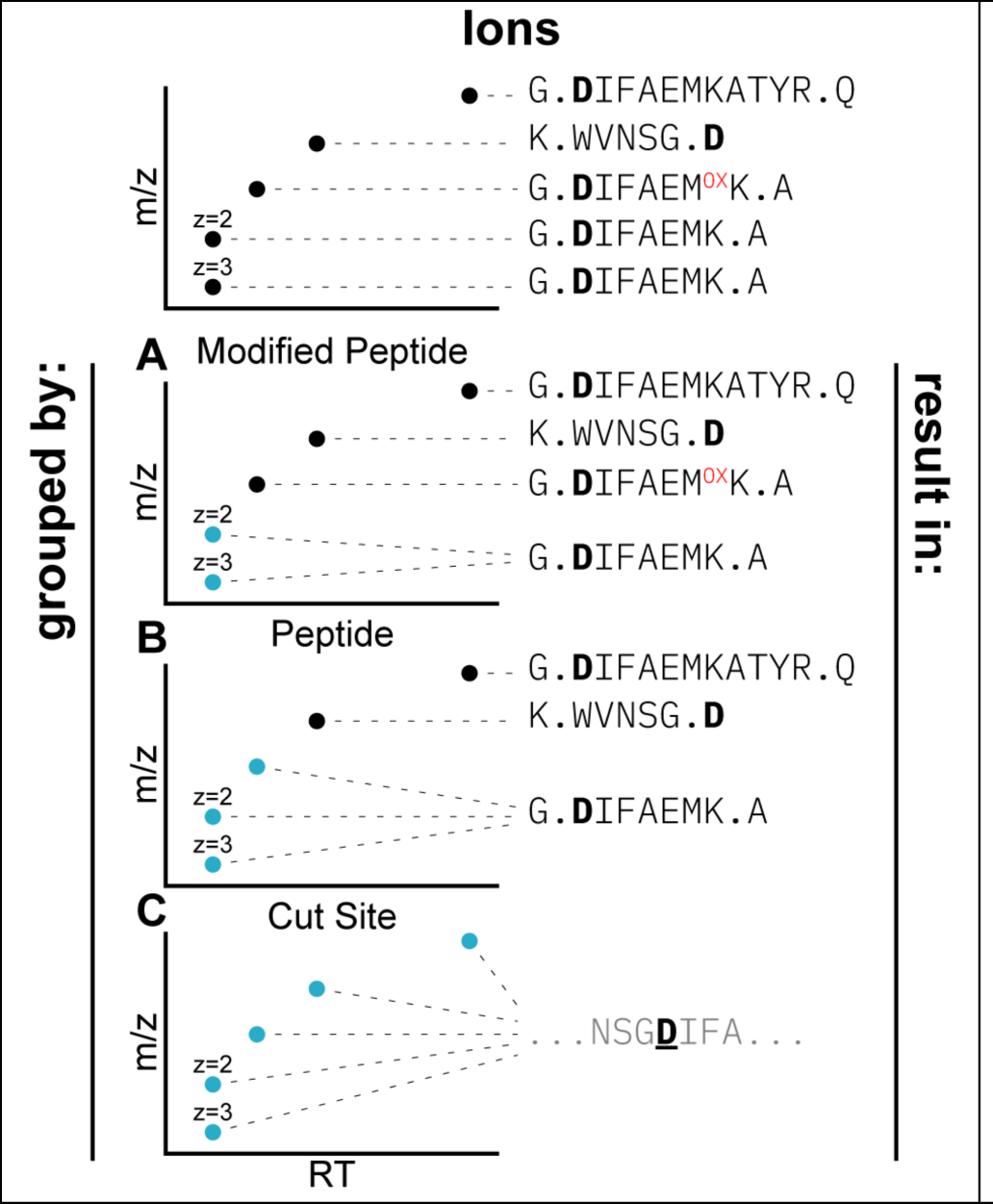
Three Merging Schemes for Ion Data. The top row represents five distinct ions that can be grouped together in three ways. (**A**) Ions that differ in charge state for the same modified peptide can be merged. (**B**) Modified peptides that differ in a specific modification (such as methionine oxidation) but correspond to the same base peptide can be merged. (**C**) Peptides that correspond to the same proteinase K cutsite can be merged. LiP-MS analyses typically merge ions to the peptide level; FLiPPR introduces further merging to the cut-site level.

Thirdly, we have found that correlating the outcome of LiP-MS refolding experiments with biophysical and structural features (such as percent disorder, isoelectric point, and domain structure) has helped illuminate key trends and have so far accumulated these metadata on a case-by-case basis. However, since many of these biophysical traits can now be calculated (or predicted) from sequence alone, valuable metadata can be generated in an automated manner, which we anticipate will be constructive for scaling up the interpretation of structural proteomics across species and clades.

Finally, FLiPPR seamlessly accepts output from FragPipe (Figure 1A), an open software suite developed by the Nesvizhskii lab that has fast, powerful, and state-of-the-art algorithms for spectral search (MSFragger)^26,27^ and LFQ (IonQuant).^26,28^ Most DDA-based LiP-MS studies to date (including ours) have employed Proteome Discoverer^29^ to perform search and LFQ (via the Minora feature detector node), though we have found that this workflow has been rate-limiting, particularly for studies that cross-compare an extensive set of conditions (e.g., LFQs with >9 raw files) which run very slowly. Indeed, our initial motivation for developing FLiPPR was to facilitate a shift to FragPipe to analyze LiP-MS data. In addition, FLiPPR creates a pipeline that formalizes some of the subtle data analysis considerations we have made in our years of experience working with LiP-MS data. Hence, we expect FLiPPR will contribute to standardizing the analysis workflow for this emerging and exciting proteomic technique by building off a popular, free platform.

## COMPUTATIONAL SECTION

### Inputs

The central input to FLiPPR is the “combined_ion” file generated by an LFQ in FragPipe using IonQuant (Figure 1A). Modifications to the default parameters are provided in Table 1, and their rationale is described in greater depth in the results section. For users less experienced with FragPipe, workflow files providing a template to specify a job are provided as supplements to this paper. Successful execution of a job generates a combined_ion file. The experiment should (minimally) consist of 6 separate raw files comprising triplicates (ideally, biological) of at least two conditions (a control and a test condition, e.g., native protein extract and refolded protein extract in a global refolding assay). Experimental designs can sometimes involve a single test condition or multiple test conditions that are all compared to a common control condition. FLiPPR can process both situations; in either case, LFQs in FragPipe are calculated with all raw files and submitted with the sequence and naming convention shown in Figure S1.

**Table 1.** Modifications to Default FragPipe Settings for LiP-MS.

| Tab<sup>a</sup> | Setting | Default Value | Value for LiP-MS |
| --- | --- | --- | --- |
| MSFragger | Precursor mass tolerance | -20–20 ppm | -10–10 ppm |
| MSFragger | Cleavage | ENZYMATIC | SEMI |
| MSFragger | Peptide mass range | 500–5000 | 500–8000 |
| Quant MS1 | MBR ion FDR | 0.01 | 0.05 |
| Quant MS1 | MBR top runs | 10 | 100<sup>b</sup> |

<sup>a</sup> Denotes which workflow tab when setting up a job in the FragPipe GUI. <sup>b</sup> Studies that include more than 10 individually collected MS files will benefit from allowing match-between-runs across all files. Ideally, this value is set greater than the number of MS files as recommended by the LFQ-MBR FragPipe tutorial.

LiP-MS studies are frequently conducted as two parallel experiments: the “LiP experiment” – which consists of all conditions including the control subjected to proteinase K and then trypsin – as well as a parallel “trypsin-only (TrP) experiment” (also called the normalization experiment) in which the samples are treated identically but the proteinase K treatment is withheld. The trypsin-only experiment is analyzed as a “standard” quantitative proteomics experiment that operates at the protein-level and addresses whether protein abundances have changed between the control and test conditions^9,10^. This second quantitative calculation is essential because abundance differences between half-tryptic peptides in the LiP experiment convolve changes in proteolytic susceptibility with changes in overall protein abundance, an effect that must be controlled. If there is a significant change in protein abundance between a test and control condition, peptide fold-changes are normalized. Hence, the second input to FLiPPR is the “combined_protein” file generated by FragPipe for the TrP experiment.

For experimental designs involving multiple tests, FLiPPR offers two ways to carry out normalization: either a single “representative” test can be compared to the control to generate a standard set of normalization factors for all tests, or alternatively, the TrP experiment can comprise all the same test conditions as the LiP experiment (giving each test/control comparison a unique set of normalization factors). Figure S1 illustrates how these two normalization schemes can be implemented.

### Processing Ion Quantifications

The combined_ion file provides raw integrated ion counts for each precursor ion (that was confidently sequenced at the MS2-level) in each of the separate runs, applying an FDR-based validation criteria to assign an ion count in runs where the peak was not identified but could be confidently “matched between runs.” At this level, FLiPPR applies a particular heuristic to managing missing data. The heuristic is applied to the set of 2*n* raw files (where *n* is the number of replicates per condition that were conducted) corresponding to each pair- wise comparison between a test condition and the control (Figure 1C). If all 2*n* ion counts (quans) are available, a simple ratio of averages is tabulated, and a P-value is calculated using a two-tailed t-test with Welch’s correction for unequal population variances (Figure 1C, case *i*). If one ion count is missing, a ratio is still calculated, but rather than treating the missing value as a zero (as commonly employed), it is dropped, and the ratio/P-value is calculated using one fewer replicate for either the test or control condition (Figure 1C, case *ii*).

A special case arises if *all* the ion counts are missing for all the replicates of one condition (either test or control; Figure 1C, case *iii*). This can be interpreted as arising from a scenario where a portion of the protein is entirely inaccessible to Proteinase K in the control condition and then becomes accessible in the test condition (or vice versa). In our experience of analyzing protein folding experiments with LiP-MS, these situations carry among the richest information since they report on large structural changes.^18,23^ However, because arguing for an effect based on missing data is fraught, a safeguard is employed that the ion must be observed in *all* replicates of the other condition (either control or test). We refer to these as “all-or-nothing” ions (Figure 1B, case *iii*). Any other combination of missing data (either because there are two missing ion counts or because one condition is missing all data, and the other condition is missing even one ion count) results in the ion being discarded and not used for quantification for this test/control comparison. For all-or-nothing ions, a ratio is calculated as a ratio of averages after Gaussian imputation for the three missing values (selected from a normal distribution with a standard deviation of 10^3^ and a mean of 10^4^ – an approximation for the limit of detection of a feature in a high-resolution MS1 scan on Orbitrap instruments). The P-value is calculated by a one-tailed Welch’s t-test. At the end of this step, a (ratio, P-value) pair is assigned to each “valid” ion for each pair-wise test/control comparison. Ions with too many missing data are discarded.

### Merging Data from Ions to Cut-Site

Most proteomics experiments seek to measure quantitative differences in protein abundance across conditions, but LiP-MS seeks to measure quantitative differences in proteolytic susceptibility at a particular cut-site. Consequently, there are several distinct ions that can be combined and averaged from the raw ion count level to the cut-site level (Figure 2). These correspond to all the ions that map to a given modified peptide (e.g., charge state 2+ or 3+), all the modified peptides that map to a given peptide (e.g., oxidized-methionine or not), and all the peptides that map to the same cut-site (e.g., the peptides [G].D^104^IFAEMKATYR^114^.[Q] and [G].D^104^IFAEMK^110^.[A] both report on the activity of proteinase K between G103 and D104 because they differ in regard to whether the subsequent trypsin digest missed the cleavage at K110). An alternative scenario where multiple peptides can encode the same cut-site arises when both half-tryptic peptides created by proteinase K are sequenced (e.g., [G].D^104^IFAEMK^110^.[A] and [K].W^99^VNSG^103^.[D]).

For all the ions that can inform on the susceptibility at a given cut-site, the (ratio, P-value) pairs for those ions are collectively considered. If they all *agree in direction* (i.e., the signs of the t-test statistics are all the same), then the ratios are combined by taking the median, and the P-values are combined with Fisher’s method to provide an updated (ratio, P-value) for the cut-site. If there are two independent ions and they disagree (e.g., the ion is more abundant in the test condition in the 2+ charge state but more abundant in the control condition in the 3+ charge state), then a median is still calculated, but the P-value is set to 1, implying there is no confidence as to whether this cut-site was more susceptible in the test or control condition. These cut-sites are discounted from the tally of the total valid cut-sites. If there are three ions, then a “majority rules” heuristic is applied: the disagreeing ion is disregarded, and the (ratio, P-value)s are only combined for the agreeing ions. In practice, it is relatively rare for more than three ions to be mapped to the same cut-site, but where this occurs, if a majority (or all) of the ions agree in direction, they are combined. If there is a tie, the P-value is set to 1.

In experimental designs with multiple test conditions, this compression is carried out separately for each test-to-control comparison. As shown in Figure 2, FLiPPR implements merging at all three levels (e.g., merge all ions together that map to the same modified peptide, peptide, or cut-site). The lowest level of merging (to a modified peptide) could be useful for studies focusing on PTMs and their effect on protein structure. The middle level of merging (to a peptide) is the one that historically we have used,^18,23^ as well as others.^8–10,20–22^ The highest level of merging (to cut-site), to the best of our knowledge, is novel to this analysis workflow.

### Output

FLiPPR produces four key output files for each test-to-control comparison: An ions file, a modified_peptides file, a peptides file, a cut-sites file, and a protein_summary file. Each row in the cut-sites file corresponds to a cut-site that was quantified by the LiP-MS experiment. A cut-site is indexed either by the host protein’s Uniprot code and the site of the PK cleavage site (for half-tryptic peptides) or by a Uniprot code and the residue range of the sequenced peptide (for tryptic peptides).

Under optimal conditions, LiP-MS experiments typically produce similar numbers of half-tryptic and full-tryptic peptides. In our experience, both kinds of peptides encode equally useful structural information, though half-tryptic peptides do provide higher structural resolution, down to the single residue-level. In contrast, tryptic peptides correspond to the *absence* of a PK cut site. They still provide quantitative information though at a lower structural resolution, and correspond to a residue range rather than an individual residue. We have generally weighted both datatypes equally, although FLiPPR output files have a column that delineates which modified peptides/peptides/cut-sites are half-tryptic, and the user can opt to consider those exclusively if desired.

Abundance ratios for each cut-site (as well as normalized abundance ratios, based on the outcome for the corresponding protein in the trypsin-only experiment) are provided along with raw P-values and FDR corrected P-values by the Benjami-Hochberg (BH) procedure for multiple hypothesis testing.^30^ BH correction is applied on a *per-protein* basis, meaning that the set of P-values for all quantified cut-sites for a given protein are subjected to FDR correction, and this process is iterated separately (and independently) for each protein. The logic for applying multiple hypothesis correction in this way is the following: Our null hypothesis is that a protein is *not* structurally perturbed by the treatment in the test condition relative to the control. Each quantified cut-site provides an opportunity to reject this hypothesis. However, the more cut-sites that are quantified for a given protein, the higher the likelihood that one of them will show a significant effect due to chance. Hence, BH correction ensures that proteins are not artificially easier to call structurally altered simply by having higher coverage.

The peptides and modified_peptides files are formatted identically, except ratios, normalized ratios, P-values, and BH-adjusted P-values are reported for each peptide or modified peptide, which involves fewer ions being merged; hence, these files contain more entries. For completeness, an ions file is provided as well (with no merging), which is like the output originally generated by FragPipe and differs solely in that it incorporates our heuristics for treating missing data.

The protein_summary file provides a high-level view of which proteins were structurally altered by the treatment and which ones were not. To do this, it counts the total number of cut-sites (or peptides) that were quantified and how many were significant according to a P-value and effect-size cutoff. We generally use 2-fold as an effect-size cutoff (|log_!_ fold– change| > 1) and 0.01 as a P-value cutoff (or alternatively, 0.05 for BH-adjusted P-values). For the largest effect sizes (|log_!_ fold– change| > 6), we relax the P-value cutoff slightly of 0.016. The user can decide whether to make the assessment at the level of peptides or cut-sites and whether to use adjusted for normal P-values. Our current recommendation is to use adjusted P-values at the cut-site level. To “call” a protein structurally altered, we typically require that two or more cut-sites (or peptides) be significant.

### FLiPPR Add-Ons: Metadata Integration

When provided with the same fasta file used as the search database during the LFQ analysis, FLiPPR can supplement the protein summary data with an array of metadata using open-source packages available in Python. Mass, molecular weight, length, and pI are obtained using Biopython.^31^ Protein disorder is predicted using Metapredict^32^ with the default disorder score thresholds. Optionally, protein domain information can be supplemented from DomainMapper,^33^ though this requires that a user provide a DomainMapper output.

Moreover, when provided with DomainMapper outputs, FLiPPR can perform advanced structural analyses by mapping cut-sites to regions within domains or linkers.

### Installing FLiPPR

Readers interested in using FLiPPR should clone the repository from GitHub at https://github.com/FriedLabJHU/FragPipe-Limited-Proteolysis-Processor. FLiPPR is built in Python and all its dependencies (pandas, scipy, numpy, metapredict, biopython, protfasta, seaborns, and matplotlib) can be installed through Python distributions and package managers.

## RESULTS

### Optimizing FragPipe LFQ for Analyzing LiP-MS Data

FragPipe features a sophisticated label-free quantification (LFQ) algorithm called IonQuant,^28^ which integrates ion intensities and performs an FDR-controlled match between runs (MBR) to include ion intensities from unassigned features. FragPipe, by default, uses a restrictive FDR cutoff (1%), which we found was less ideal for LiP-MS experiments, particularly in the context of our heuristics for managing missing data (Figure 3A).

**Figure 3.**
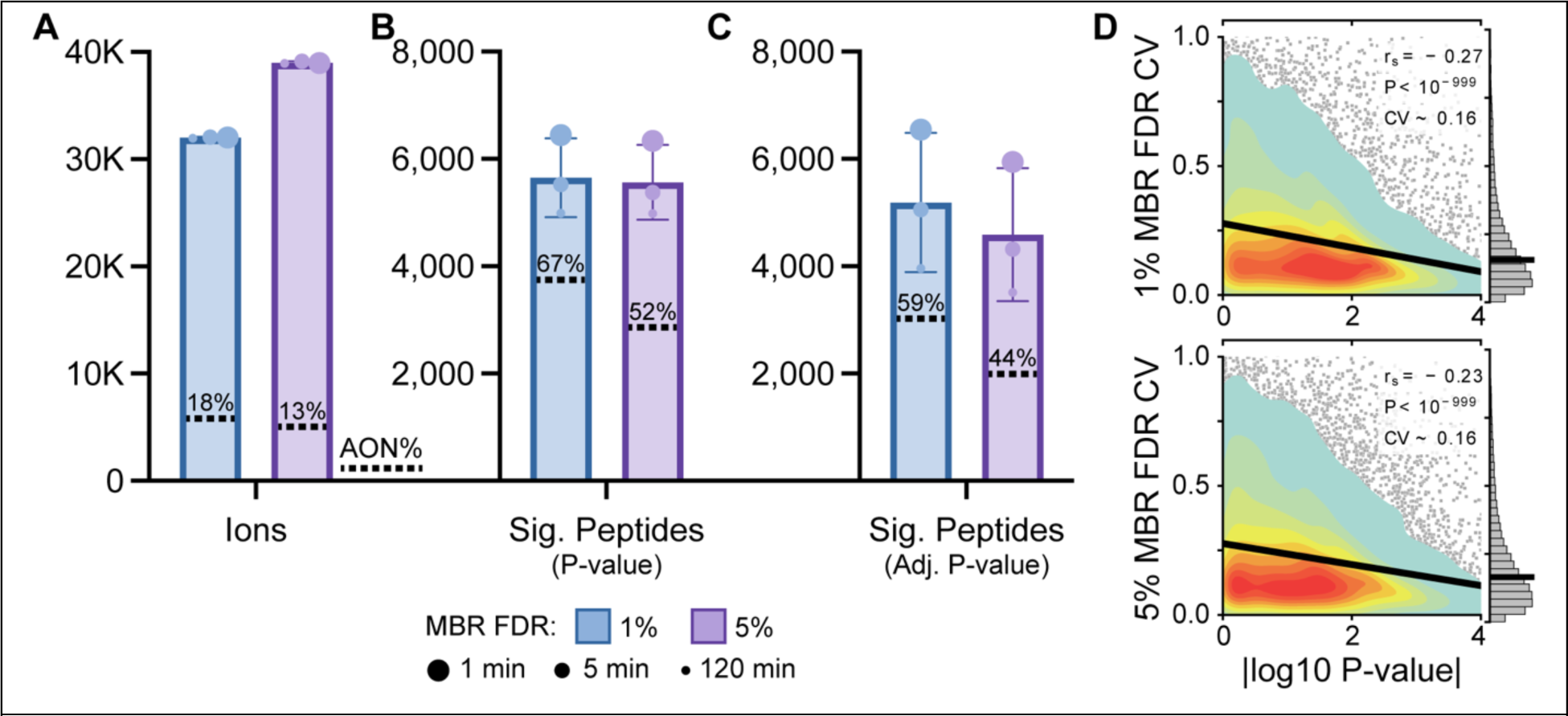
The match-between-run (MBR) false discovery rate (FDR) is a key parameter for quantifying LiP-MS data. (**A**) The total number of quantified ions in an experiment in which *E. coli* extracts were globally unfolded and refolded for 1, 5, or 120 min before performing limited proteolysis, with the default (1%) or adjusted (5%) MBR FDR parameter. Data were processed in FLiPPR with 12 total raw files (3 replicates of a native extract, used as a control, and 3 replicates each of the 3 different refolding times, creating 3 different test conditions). The percent of ions that were all-or-nothing (AON) is marked. (**B**) Following the merging of ions into peptides, the number of peptides that are significantly different between refolded and native (P < 0.01; or P < 0.016 for effect-sizes larger than 64-fold), and what proportion of them were AON. (**C**) As panel B, except using Benjami-Hochberg adjusted P-values instead (the threshold for significance is Adj. P < 0.05). (**D**) Quality control plots for the two MBR FDR settings, showing the coefficient of variation (CV) for each quantified peptide in the refolding reaction (shown for the 1 min refolding time) as a function of the P-value that it is distinct from the native sample. Insets provide Spearman’s rank correlation coefficient (*rs*), P-value, and median CV across all quantified peptides.

Missing values have an outsized significance in LiP-MS experiments since half-tryptic peptides that only appear in the test (or control) samples nominally report on the most profound structural changes since they imply a region was inaccessible to PK in one condition and became accessible in the other. When we raise the FDR cutoff from 1% to 5%, we find that the number of quantified ions goes up considerably, but importantly, the number of all-or-nothing (AON) ions *decreases* (Figure 3A). We considered this change because we are quite restrictive in how much missing data causes an ion to become excluded, so we thought it advantageous to initially be more permissive toward which features are mapped to an ion. At the same time, by raising MBR FDR, we are making it *easier* to include a feature into an ion’s quantification set, which in turn makes it *harder* for data to be missing and, in turn, *harder* for an ion to be retained as an AON. The net result is that by being more inclusive, we are more confident when an ion is truly an AON.

This precaution is merited because even though AON ions are rare, they make up a disproportionate fraction of the peptides that rise to the level of being significantly different between conditions (Figure 3B). Higher MBR FDR lowers the percentage of significant peptides that are AON, leaving behind the ones that – we believe – are more confident. BH-correction to P-values culls the number of AON peptides that are labeled significant (Figure 3C) at a similar frequency to the other significant peptide; this is because although AON peptides possess the largest effect-sizes, they also tend to have greater variability in the replicates of the condition where they are observed.

Of course, the potential cost to raising the MBR FDR is that more unassigned features’ ion intensities will be spuriously grouped into an ion quantification set to which they do not belong. This would then be reflected in ion abundances becoming noisier, resulting in larger coefficients of variation (CVs) across replicates. In practice, this does not occur, and the quality control plots (shown in Figure 3D) reveal that the CV distribution and median are essentially unchanged even with this 5-fold increase in the MBR FDR cutoff. We surmise that the reason this is the case is that we discard ions with even two missing values, so features in two separate replicates would have to be spuriously matched before the ion would be used for quantification.

FLiPPR automatically generates plots like the ones shown in Figure 3D, which we have employed as a basic data quality control metric. These charts show, for each quantified peptide, its CV in the test condition (i.e., refolded) – which charts the amount of reproducibility in the refolding reaction and in the limited proteolysis at that site – as well as the P-value against the null hypothesis that the peptide is present in different abundances between the test and control. A “healthy” sample should have a median CV between 10-20%, a majority of the peptides not rejecting the null hypothesis (|log10 P-value| < 2), and a negative trend between the two (less noisy data are more likely to reject null hypotheses).

### Merging Peptides into Cut-sites Improves Statistical Confidence

Figure 4A shows a representative peptide-level volcano plot from one of our LiP-MS experiments refolding the proteome of *E. coli*. Each point represents a confidently identified and quantified peptide. We typically find that half-tryptic peptides are more abundant in refolded samples compared to native (Figure 4B), a characteristic that is qualitatively consistent with the notion that misfolded proteins are less well-packed and are, therefore, more susceptible to PK than natively folded proteins. In general, we find this feature any time the test condition is one that “perturbs” proteins and have seen it in a number of ongoing studies focusing on other variables. Following the fate of the ca. 32,500 peptides quantified in this experiment (Figure 4C), we find that a relative minority (602) are discounted due to the ions that merge into it showing inconsistent signals.

**Figure 4.**
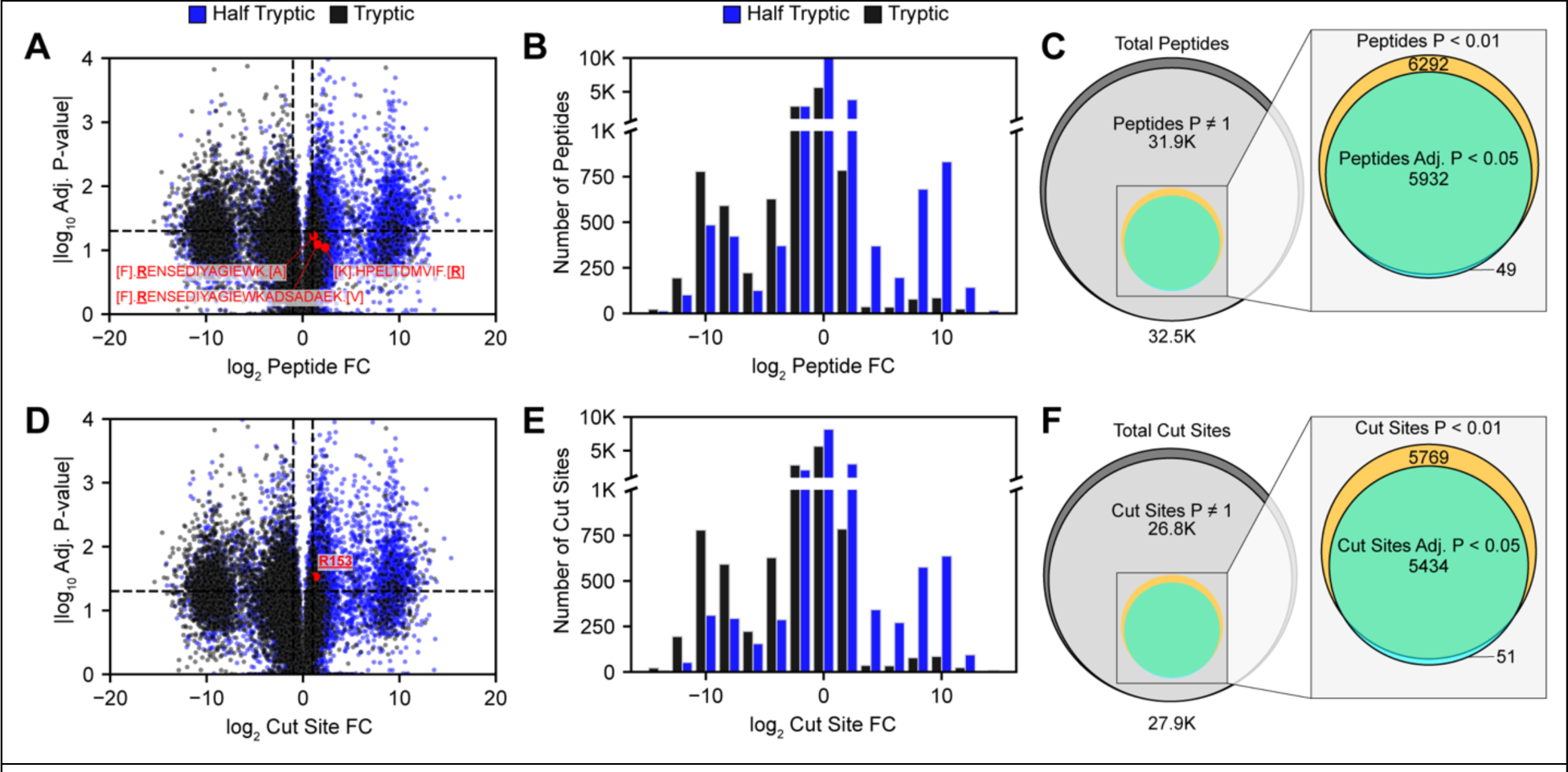
Analysis of LiP-MS at the Peptide or Cut-site Level. (**A**) Representative volcano plot in which each dot represents a peptide quantified in the experiment. Three specific peptides are highlighted in red. Data are from the 1 min time point from refolding assays on *E. coli* extracts and reflect the abundance ratio of refolded/native. Blue corresponds to half-tryptic peptides, and black to tryptic peptides. (**B**) Histogram of peptide abundance ratios. Enrichment of half-tryptic all-or-nothings in the refolded form is frequently encountered after global refolding or other test conditions that result in perturbed structures. (**C**) Accounting of the peptides. Dark gray = all peptides with sufficient data to qualify; light gray = all “valid” peptides (e.g., discounting those with inconsistent ions); mustard = significant peptides based on effect-size (>2-fold) and P-value (< 0.01, or <0.016 if effect-size > 64); teal = significant peptides based on effect-size (>2-fold) and BH-adjusted P-value (< 0.05). (**D-F**) As panels **A-C,** except at the cut-site level rather than the peptide level. The cut-site at F341 in gpmI that results from the merger of the three peptides shown in **A** is highlighted in red in panel **D**.

FLiPPR keeps these “invalidated” peptides in the output for completeness but sets their P-value to 1. A minority of these peptides are deemed significant (mustard circle) by the cutoffs required to call it structurally altered, and these are further culled to a smaller set by FDR correction (teal circle).

FLiPPR introduces the idea of merging together peptides that map to the same PK cut-site. Following this process, we redraw the volcano plot and relative abundance histogram (Figure 4D-E), where the points represent cut-sites instead of peptides. One example, highlighted in red, shows a scenario in which three separate peptides, none of which were statistically significant on their own, admitted a significant cut-site once the ions were merged by Fisher’s method. In this LiP-MS experiment, there are 27,900 cut-sites (instead of 32,500 peptides), but 5,769 cut-sites are assessed as significant (Figure 4F; instead of 6,292 peptides). This slight reduction arises primarily from “removing” duplicates, but it is noteworthy that only 8% of significant peptides are lost even though 14% of the peptides were merged. This difference occurs because some peptides cross the threshold to significance when their data are merged. Hence, we assess that using cut-sites instead of peptides results in a dataset with fewer duplications and more “unique” significant sites detected.

### Protein-level Trends in LiP-MS Datasets become Sharper and More Confident with FLiPPR Analysis

Our original work on refolding the *E. coli* proteome highlighted the prevalence of nonrefoldability amongst soluble proteins^18^, and estimated that after 1 min of refolding, 56% of *E. coli* proteins could return to native-like structures, a figure that rises to 67% after providing 2 h to refold. To et al. also called attention to the fact that differences in protein coverage are a source of bias: Proteins with more identified peptides are more likely to be labeled as “structurally altered” because there are more opportunities for a significant effect to be detected.

We sought to compare how these overall outcomes of the experiment are affected by some of the differences in the analysis implemented in FLiPPR (Figure 5A- C). If we analyze the data in a manner analogously to the original work (Figure 5A), we find that refoldability levels go down (48% at 1 min and 53% at 2 h). This difference is in part due to coverage bias, arising because FragPipe’s search (with MSFragger) produces more identifications than ProteomeDiscoverer v2.3 (with Sequest) does: Namely, 31,900 and 31,800 peptides at 1 min and 2 h respectively versus 28,700 and 28,200 in ProteomeDiscoverer. As mentioned, proteins with more identifications are “easier” to label as structurally altered, which can be corrected by adjusting for multiple hypothesis testing. Using adjusted P-values (Figure 5B), the refolding propensities become closer to originally reported values (55% at 1 min and 66% at 2 h), and these overall trends are not appreciably changed if cut-sites are used instead of peptides (Figure 5C).

**Figure 5.**
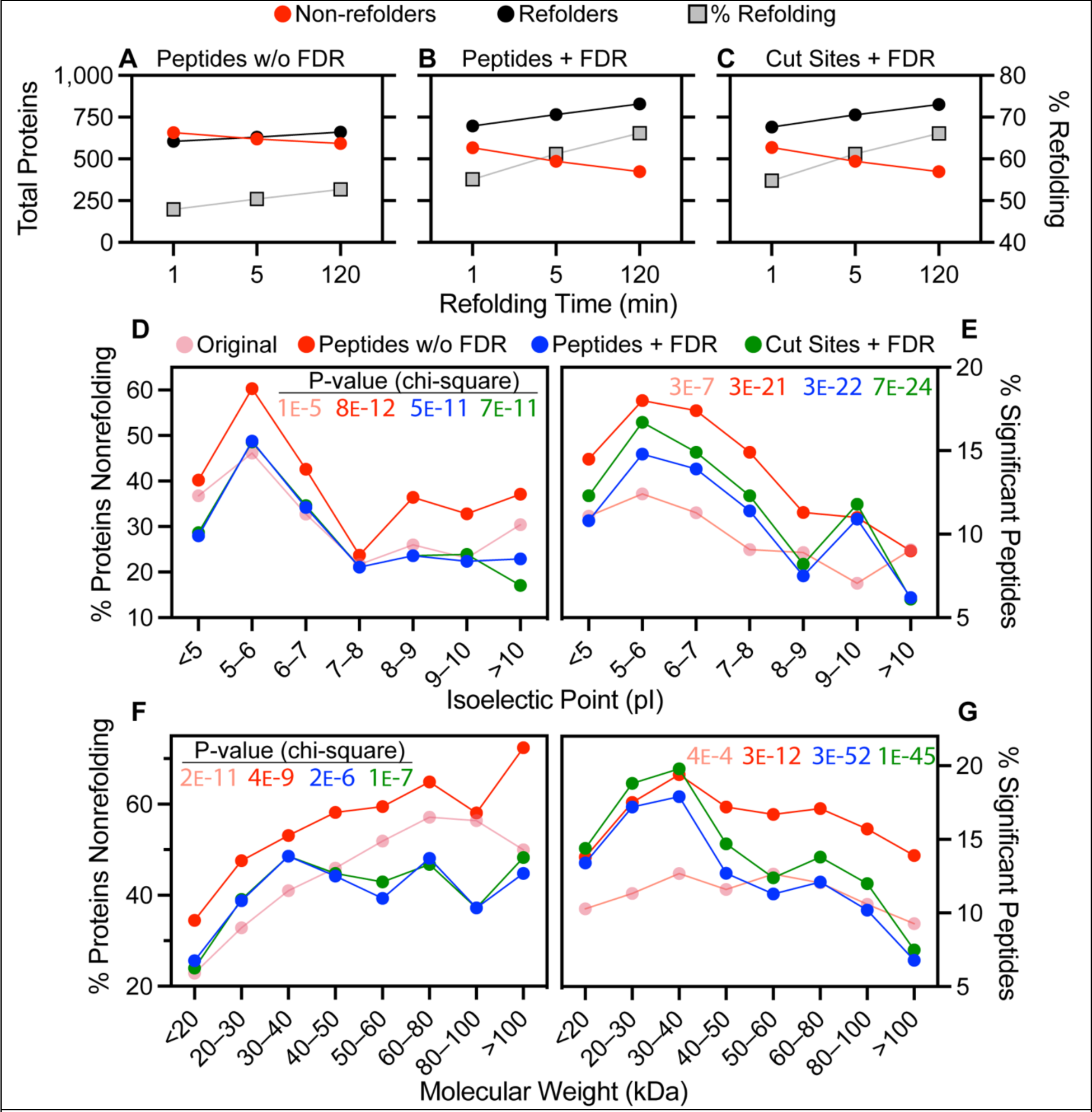
Reanalysis of Refoldability Data with FLiPPR Sharpens Trends. (**A-C**) Plots show the number of proteins (left y-axis) labeled refoldable (black) and nonrefoldable (red) and the fractional refoldability (right y-axis, gray boxes) as a function of refolding time, depending on whether calls are made based on peptides without Benjami-Hochberg FDR correction (**A**), with FDR correction (**B**), or with FDR correction and after merging peptides to cut-sites (**C**). Data correspond to *E. coli* refolding assays from ref. 18 (PRIDE: PXD025926). (**D**) Fraction of proteins nonrefolding as a function of protein isoelectric point (pI), based on the original analysis (pink) or using FLiPPR using one of three schemes. Inset shows P-value from chi-square test against null hypothesis pI does not explain differences in refoldability. I Fraction of peptides (or cut-sites for green symbols) that are assessed as significantly different after refolding, as a function of the pI of the protein from which they came, as based on the original analysis (pink) or using FLiPPR using one of three schemes. Inset shows P-value from chi-square test against null hypothesis pI does not explain differences in refoldability. (**F**) As **D** except proteins divided by molecular weight. (**G**) As **E** except peptides divided by the molecular weight of the protein from which they came.

Global protein refolding assays also demonstrated that refoldability possessed clear correlations with other biophysical and biochemical variables. For instance, from all the studies we have conducted to date, a general theme has emerged that proteins at the extrema of the pI range (the most acidic and the most basic) refold more often than those whose pI are between 5–6 (mildly acidic proteins). Impressively, the pI trend become *much* sharper when the exact same data are analyzed in FLiPPR, taking advantage of the two key improvements we have implemented. Based on the 5-min timepoint, the original analysis showed that proteins whose pI is between 5–6 did not refold 46% of the time, a fraction that decreases for the proteins in the <5 (37%, 0.80- fold) or >10 (30%, 0.65-fold) pI ranges. Using FLiPPR (Figure 5D, red trace), the nature of acidic and basic proteins to refold better is more striking. As before, the peak of nonrefoldability (60%) occurs at pI 5–6, but this drops off more dramatically for proteins with pI < 5 (40%, a 0.66-fold) or >10 (37%, a 0.62-fold). The slopes become even steeper after FDR correction has been incorporated (Figure 5D, blue trace). Now, pI < 5 (pI > 10) proteins have nonrefoldability rates that are 0.57-fold (0.47-fold) that of the maximum. If we accept as a ground-truth that acidic and basic proteins are more refoldable than those with pI 5–6 (as we have seen in all our studies to date),^18,23^ then this implies that analyzing LiP-MS data with FLiPPR provides a view that coheres much more closely to physical reality.

Another metric that shows the FLiPPR-analyzed dataset has more discriminating power is the chi-square test on the null hypothesis that pI does *not* explain differences in protein refoldability. In the original publication, the P-value was placed at 1.3 × 10^-5^; with FLiPPR it becomes closer to 10^-11^ (depending on whether peptides or cut-sites are used). We emphasize here that this is purely from reanalyzing the exact same raw mass spectra.

One of the signs that the pI-refoldability trend is robust is that it is apparent at both the protein level (i.e., assessing proteins as refoldable or not after grouping peptides/cut-sites by protein) and at the individual peptide/cut-site level (i.e., not grouping these by protein and simply calculating the percent that are significant across categories). At the peptide/cut-site level, we find that FLiPPR produces much sharper trends than the original analysis (Figure 5E), and the trends become even more apparent as we introduce FDR correction and peptide to cut-site merging. The strengthening of the correlation is apparent in the chi-square tests as well.

The original work studying refoldability of *E. coli* proteins documented an apparent trend whereby proteins of greater molecular weight were more likely not to refold (Figure 5F, pink trace). However, it was noted that this trend could be the result of coverage bias. In general, the peptides assigned to larger proteins were not more likely to be significant (Figure 5G, pink trace), though larger proteins do generally get more peptides mapped to them, thereby making it more likely that one of them would be significant. Reexamining this trend is therefore a stringent test for whether our FDR correction can “catch” this problem. Indeed, using FLiPPR with FDR corrections (blue and green traces in Figure 5F), we find that there is *some* correlation between molecular weight and refoldability upto 40 kDa, but afterwards, the trend is flat. This is very well recapitulated at the peptide/cut-site level (Figure 5G), in which peptides are indeed more likely to be significant in 30–40 kDa proteins than in proteins <20 kDa, but then the trend reverts. This finding makes sense because 30–40 kDa is the size of the largest single domains and our previous findings have found that larger domains typically refold poorly; proteins larger than this typically contain multiple domains.

Hence, we conclude that the more reliable and confident quantification offered by FLiPPR improves downstream analysis of LiP-MS data, and in the case of the present renalaysis of our original study, firms up one finding (pI) and offers a reinterpretation of another (molecular weight).

### FLiPPR Enables Rapid Analysis of Data from Non-Model Species, Facilitating Cross-Species Comparisons

Work in our laboratory has focused on using LiP-MS as a tool to interrogate protein folding on the proteome-scale,^18,23^ with a focus on addressing how “refoldable” are proteins; namely, if they are globally unfolded in 6 M guanidinium chloride and compelled to refold by dilution, how many proteins succeed in this challenge and can assume a conformation identical to their native ones that were never unfolded? We have found this can be assessed in a high-throughput manner through LiP-MS, in which the proteolysis pattern of a native extract and one that is subjected to an unfolding/refolding cycle are compared. A current goal of ours is to perform these assays on many organisms to search for evolutionary trends in refoldability, which in turn prompted us to build computational workflows with higher speed and reliability. For this study, we have refolded the proteome of *Bacillus cereus (ATCC 14579)*^34^ following an approach similar to those previously described.^18,23^ The methods are described in full in the Experimental Section. In brief, we grew triplicate cultures of *B. cereus* to a final OD600 of 1.0, lysed by cryogenic pulverization, subjected the lysates to global unfolding and refolding, and performed limited proteolysis on the native extract and the refolded ones. Refolding was monitored at three time points following dilution: 1 min, 5 min, and 30 min.

In previous studies, acquisition of biochemical and biophysical metadata for each protein provided an important set of metadata which enabled correlates with refoldability to be determined from LiP-MS experiments. However, we relied on databases (EcoCyc^35^ for E. coli, and SGD^36^ for yeast), which are not available in general for non-model organisms. To make metadata acquisition more streamlined and species-agnostic, we have implemented in FLiPPR a basic infrastructure to automate the acquisition of metadata (Figure 6A) using BioPython prediction tools and UniProt.

**Figure 6.**
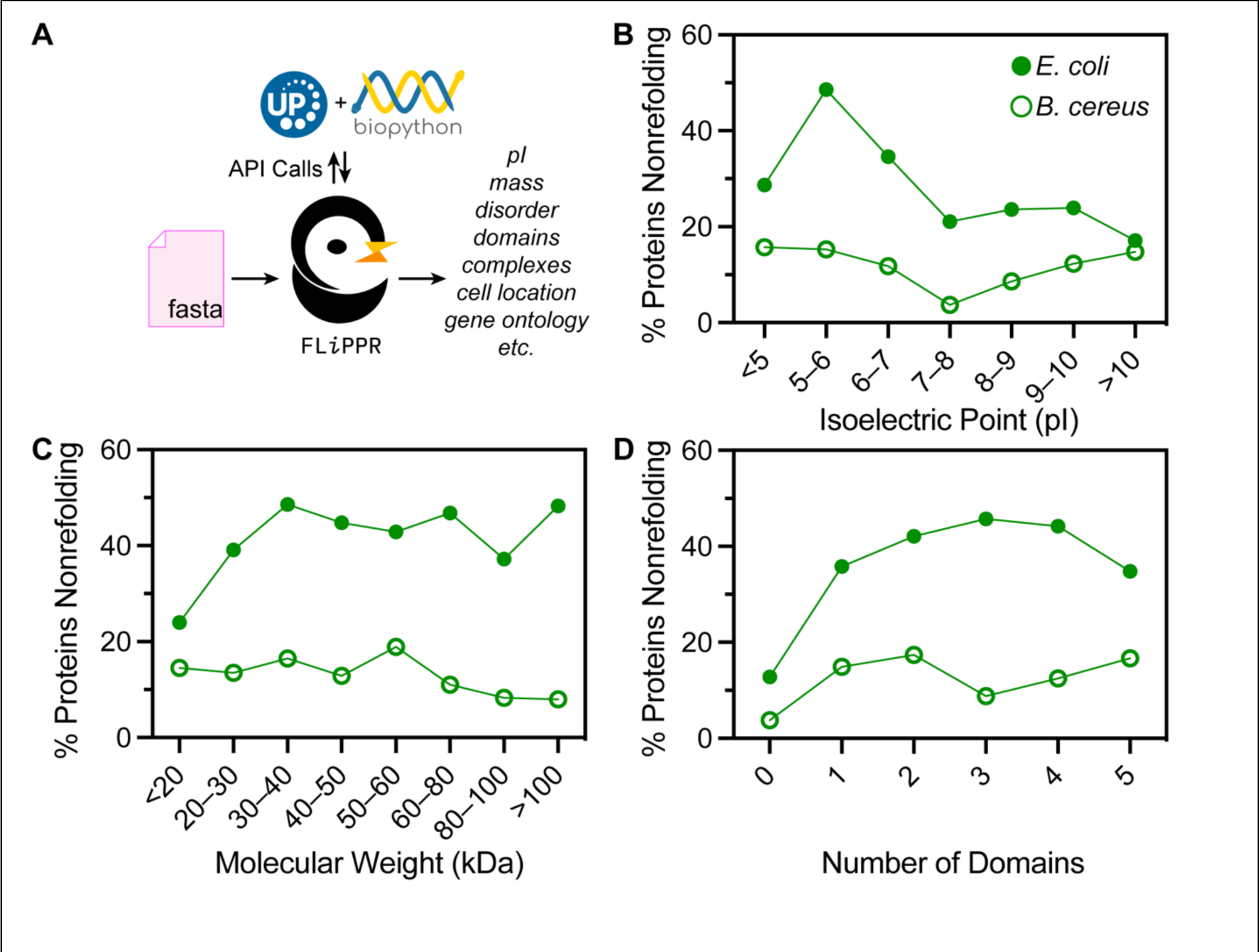
Comparison of nonrefolding rates between two species. (**A**) Scheme illustrating how protein metadata is retrieved within FLiPPR. The only required input is the same FASTA file provided to FragPipe during the primary search and quantification. FLiPPR produces a metadata .tsv file for the whole proteome, whose information is then merged into the output files described above. (**B**) Fraction of proteins nonrefolding as a function of protein isoelectric point (pI), based on cut-sites with FDR correction for two organisms, *E. coli* (ref. 18) and *Bacillus cereus* (data reported in this study). pI is calculated in BioPython. (**C**) Fraction of proteins nonrefolding as a function of molecular weight (MW), based on cut-sites with FDR correction for *E. coli* and *B. cereus*. MW is calculated in BioPython. (**D**) Fraction of proteins nonrefolding as a function of number of domains, based on cut-sites with FDR correction for *E. coli* and *B. cereus*. Domain assignment is determined in DomainMapper. 37

When we analyzed the refolding LiP-MS data from *B. cereus* in FLiPPR, it was clear that although its proteins refold on the whole more efficiently than *E. coli’s*, several trends are conserved between the two organisms whilst others are different. For instance, we find in *B. cereus* that mildly acidic proteins (pI 5–6) are still amongst the worst refolders (like *E. coli*), although one major point of difference is that proteins at the extremes of the pI range (<5, >10) are also relatively poor refolders, a behavior quite distinct from *E. coli* (Figure 6B). Like *E. coli*, we find a relatively flat dependence on molecular weight for proteins larger than 30 kDa (Figure 6C). Moreover in both *E. coli* and *B. cereus*, we find that disordered proteins (which contain no globular domains) refold the best, and there is a “shallow” additional challenge to refold as the number of domains increases beyond one (Figure 6D). Whether or not bacterial proteomes generally refold quite poorly (like *E. coli*) or quite efficiently (like *B. cereus*) is a question of current active research.

## DISCUSSION

In recent years, FragPipe has become recognized as a leading proteomics analysis platform which combines a fast and sensitive search engine and several quantification algorithms (with support for LFQ, SILAC, and TMT). It is also free, easy to use, and supports both data-dependent acquisition (DDA) and data-independent acquisition (DIA) modalities. Structural proteomics – an emerging field within proteomics – currently suffers from having a panoply of software pipelines which hampers interoperability and standardization. For instance, there are at least 10 different packages for analyzing crosslinking mass spectrometry data.^15^ LiP-MS studies have historically employed primarily ProteomeDiscoverer (for DDA) and Spectronaut (for DIA) with less standardization in how raw quantifications are processed. Likewise, FPOP data has historically been analyzed in ProteomeDiscoverer with user-defined workflows to handle the experiment’s specific needs. Recently, a standardized workflow for FPOP experiments in FragPipe was proposed^38^ showing that this platform is well-suited to bring its numerous other strengths to bear in structural proteomics. Our original goal in developing FLiPPR was to create a pipeline that would facilitate using FragPipe to analyze LiP-MS data. Following this goal, we have aimed to build a tool that is easy to use, compatible with various experimental designs, provides a range of useful outputs (including those for quality control), and facilitates metadata integration.

LiP-MS has been applied to answer a range of biological and biophysical questions. Our lab’s focus on protein refolding – which can produce very divergent protein conformations – has made us particularly sensitive to the importance of missing data in LiP-MS, and how much useful information can be gained from them if used carefully. FLiPPR formalize these heuristics into software that can be widely used for structural proteomics. Compared to our previous work, we have incorporated two further improvements: (1) a protein-centric FDR correction to control for coverage bias; and (2) a hierarchy of data merging rules that enables quantification from the ion level to the cut-site level. These two additions create more robust and less redundant datasets. The result is greater accuracy, which can be attested by sharper trends and correlations upon reanalyzing earlier work on the *E. coli* proteome (cf. Figure 5). As we proceed with LiP-MS experiments on proteomes from more diverse organisms (such as *B. cereus*, discussed here), we expect that the speed and standardization offered by the FragPipe/FLiPPR pipeline will prove invaluable.

There are several improvements we foresee adding to future versions of FLiPPR. The Picotti lab introduced a mixed linear model to perform normalizations that considers data from LiP and TrP experiments collectively, rather than basing decisions on whether to normalize by the TrP experiment unilaterally.^10^ In practice, we have found that when LiP-MS is applied to biophysical questions (and the test and control conditions arise from the same biological source, but differentiate from each other by treatments performed *in vitro*), normalization is less critical and, in some cases, weakens the data by propagating more error to the quantifications. On the other hand, we acknowledge that careful normalization is required for some biological studies and that this option should be added to increase the generality of FLiPPR to other LiP-MS applications.

A second outstanding question is how to apply LiP-MS to interrogate the effect of post-translational modifications (PTMs) on protein structure. PTMs can profoundly impact protein structure,^39,40^ and by extension their limited proteolysis pattern, making LiP-MS a potential technique to map this effect. For now, FLiPPR provides an output that restricts merging to the modified peptide level, and users can interrogate these files to see (for instance) if a phosphorylation within a peptide results in a different outcome compared to the same unmodified peptide. In practice, allostery can induce structural changes far from the PTM-site, which creates a “proteoform problem”: a half-tryptic peptide associated with a PTM need not be close to the PTM itself, and so sequencing such a half-tryptic peptide would not provide enough information to trace it back to the specific (set of) PTM(s) that induced it. Due to this ambiguity, we recommend merging over peptide modifications, and regard LiP-MS experiments as probing the “average” of all proteoforms for a given protein. While separating peptide modifications more explicitly into proteoform-specific categories is a functionality we plan to add, we emphasize that new LiP-MS experiments (potentially of a middle-down nature) will be needed to confidently assess the effect of PTMs on structure.

In summary, we anticipate growth in the number and variety of applications for LiP-MS and hope that FLiPPR will contribute by lowering the barrier to adopt this structural proteomic approach and standardizing statistical procedures for LiP-MS data analysis.

## ASSOCIATED CONTENT

### Supporting Information

The Supporting Information is available free of charge at XXX. It includes the experimental section (for *B. cereus* refolding), and Figure S1 (describing naming conventions for how to call FLiPPR based on naming convention for raw files inputted into FragPipe).

## AUTHOR INFORMATION

### Corresponding Author

Stephen D. Fried – Department of Chemistry and T. C. Jenkins Department of Biophysics, Johns Hopkins University, Baltimore, MD 21218, United States. Orcid: https://orcid.org/0000-0003-2494-2193 ;

### Authors

Edgar Manriquez-Sandoval – T. C. Jenkins Department of Biophysics, Johns Hopkins University, Baltimore, MD 21218, United States. Orcid: https://orcid.org/0000-0001-7284-1237 ;

Joy Brewer – Department of Chemistry and Biochemistry, Old Dominion University, Norfolk, VA, 23529, United States. Orcid: https://orcid.org/0000-0003-2758-351X ;

Gabriela Lule – Department of Chemistry, Johns Hopkins University, Baltimore, MD 21218, United States. Orcid: https://orcid.org/0009-0002-9419-9526 ;

Samanta Lopez – Department of Chemistry, Johns Hopkins University, Baltimore, MD 21218, United States. Orcid: https://orcid.org/0009-0001-6768-8802 ;

### Notes

The authors declare no competing financial interest. Raw proteomic data have been submitted to PRIDE via theProteomeXchange under the accession codes XXXX.

## ACKNOWLEDGEMENTS

We would like to thank the NSF Division of Molecular and Cellular Biology for a CAREER grant (2045844) and NIH/NIGMS for a New Innovator Award (DP2- GM140926) both to S.D.F. E.M.S. thanks the Program in Molecular Biophysics training grant (NIH-T32GM135131). Funds from the NSF CAREER grant and the Dreyfus Foundation partly supported undergraduates involved in the project (J.B., G.L., S.L.). We thank Alexey Nesvizhskii for critical manuscript reading and helpful feedback.

## Supporting Information

### Culture and Lysis of Bacillus cereus (ATCC 14579)

Saturated overnight cultures of B. cereus cells (ATCC 14579) were used to inoculate 3 × 100 mL (biological triplicate) of ATCC Medium: 3 in 250 mL flasks at a starting OD600 of 0.05. Cells were incubated at 37°C with agitation (220 rpm) to a final OD600 of 1.0, followed by centrifugation at 3000 rcf for 10 min at 4°C. Supernatants were removed, cell pellets where flash frozen with liquid nitrogen for 30 seconds, and stored at –80°C until further use.

Cell pellets were resuspended in 1 mL of native buffer (20 mM Tris pH 8.2, 150 mM KCl, 2 mM MgCl2) with the addition of 20 units of DNase I (NEB M0303S) and protease inhibitors (500 µM PMSF; Thermo Scientific 36978, 15 µM E-64; Millipore Sigma E3132, 50 µM Bestatin; Millipore Sigma B8385). Resuspended cells were flash frozen by slow drip over liquid nitrogen followed by cryogenically pulverized with a freezer mill (SPEX Sample Prep) over 8 cycles consisting of 1 min of grinding, 9 Hz, and 1 min of cooling. Pulverized lysates were transferred to a 50 mL centrifuge tube and thawed at 4°C. Lysates where then transferred into 1.5 mL microcentrifuge tubes and clarified at 16,000 rcf for 15 min at 4°C to remove cellular membrane and debris. Clarified lysates were then transferred into 3 mL Beckman Coulter Konical tubes (Beckman Coulter C14307) in preparation for ultracentrifugation at 33,300 rpm at 4°C for 90 min using a SW55-Ti rotor to deplete ribosomes without a sucrose cushion. These clarified supernatants were then transferred into new 1.5 mL microcentrifuge tubes and protein concentrations were determined using a bicinchoninic acid assay (Rapid Gold BCA Assay, Pierce) in a clear 96-well plate (Corning Falcon 353075) with a plate reader (Molecular Devices iD3).

Using the results from the BCA Assay, the clarified lysates were normalized to a protein concentration of 2.0 mg mL^−1^ using the same native buffer (20 mM Tris pH 8.2, 150 mM KCl, 2 mM MgCl2). The normalized lysate is used as the starting point for all the following workflows.

### Preparation of Native and Refolded Lysates

Native lysates were prepared by diluting 58 µL of normalized lysate with native buffer supplemented with 5.7 mg of guanidinium chloride (GdmCl) and 15.4 µg of dithiothreitol (DTT) such that the final composition of the Native lysates was: 0.116 mg mL^−1^ protein, 20 mM Tris pH 8.2, 150 mM KCl, 2 mM MgCl2, 0.1 mM DTT, and 0.06 M GdmCl in a 1 mL volume. These samples were prepared in biological triplicate and allowed to incubate for at least 1 hour at 25°C prior to limited proteolysis.

Unfolded lysates were prepared by concentrating 290 µL of normalized lysate supplemented with 28.7 mg of GdmHCl and 77.1 µg of DTT in a Vacufuge Plus (Eppendorf) to a final volume of 50 µL. The final composition of the Unfolded lysates was: 11.6 mg mL^−1^ protein, 116 mM Tris pH 8.2, 870 mM KCl, 11.6 mM MgCl2, 10 mM DTT and 6 M GdmCl. These samples were prepared in biological triplicate and allowed to incubate for at least 16 hours at 25°C prior to refolding and subsequent limited proteolysis.

Refolded lysates were prepared by diluting 5 µL of Unfolded lysate into 495 µL of refolding buffer (19 mM Tris pH 8.2, 143 mM KCl, 1.9 mM MgCl2). The final composition of the Refolded lysates was: 0.116 mg mL^−1^ protein, 20 mM Tris pH 8.2, 150 mM KCl, 2 mM MgCl2, 0.1 mM DTT, and 0.06 M GdmCl in a 500 µL volume. These samples were prepared in biological triplicate and allowed to incubate for 1, 5 and 30 minutes at 25°C prior to limited proteolysis.

### Limited Proteolysis of Native and Refolded Lysates

A stock of PK solution was freshly prepared at a concentration of 0.116 mg mL^−1^ Proteinase K in native buffer with 10% glycerol. Triplicate limited proteolysis (LiP) samples of Native and Refolded conditions, for all time points, were then generated by aliquoting 200 µL of each lysate into new 1.5 mL microcentrifuge tubes containing 2 µL of PK solution (1:100 enzyme:substrate mass ratio), quickly aspirated 10 times, and incubated for exactly 1 min at 25°C. Samples were then transferred to a 110°C mineral oil bath for 5 minutes to inactivate Proteinase K. Boiled samples were then transferred into new 1.5 mL microcentrifuge tubes containing 150 mg of urea to achieve a final volume of 312 µL and 8 M urea concentration. Triplicate trypsin-only (TrP) controls of the Native and Refolded samples were generated in the same way without the addition of PK solution. This process generated the following 18 samples: 3 × Native_TrP, 3 × Refolded_TrP, 3 × Native_LiP, 3 × Refolded_1min_LiP, 3 × Refolded_5min_LiP, and 3 × Refolded_30min_LiP.

All samples then received 6.24 µL of freshly prepared 500 mM DTT (final concentration 10 mM DTT) to reduce disulfide bonds and incubated for 30 min at 37°C with agitation 700 rpm followed by 17 µL of freshly prepared 750 mM IAA (final concentration 40 mM IAA) and incubated for 45 min in the dark at 25°C to alkylate reduced cysteines. Next, 2 µL of 0.116 mg mL–1 of LysC (NEB P8109) were added to each sample and incubated for 2 hours at 37°C. Samples were then diluted with 1005 µL of 100 mM ammonium bicarbonate followed by 4 µL of 0.116 mg mL^−1^ of Trypsin-ultra (NEB P8101) and incubated at 25°C for 16–24 hours at 25°C. Trypsin was then quenched with 1% v/v trifluoroacetic acid prior to desalting with Sep-Pak C18 1cc cartridges (Waters).

Cartridges were first conditioned (2 × 1 mL 80% ACN, 0.5% TFA) and equilibrated (4 × 1 mL 0.5% TFA) before samples were slowly loaded under a weak vacuum. The columns were then washed (4 × 1 mL 0.5% TFA), and peptides were eluted by addition of 1 mL of elution buffer (80% ACN, 0.5% TFA). During elution, vacuum cartridges were suspended above 15 mL conical tubes, placed in a swing-bucket rotor (Eppendorf 5910R), and spun for 5 minutes at 300 rcf. Eluted peptides were transferred from Falcon tubes back into new 1.5 mL microcentrifuge tubes and dried using a Vacufuge Plus (Eppendorf). Dried peptides were stored at −80 °C until analysis. For analysis, samples were vigorously resuspended in 0.1% FA in Optima water (ThermoFisher) to a final concentration of 1 mg mL^−1^.

### Collecting MS^2^ Spectra

Chromatographic separation of digests was carried out on a Thermo UltiMate3000 UHPLC system with an Acclaim Pepmap RSLC, C18, 75 μm × 25 cm, 2 μm, 100 Å column. Approximately, 1 μg of protein was injected onto the column. The column temperature was maintained at 40 °C, and the flow rate was set to 300 nL min^−1^ for the duration of the run. Solvent A (0.1% FA) and Solvent B (0.1% FA in ACN) were used as the chromatography solvents.

The samples were run through the UHPLC System as follows: peptides were allowed to accumulate onto the trap column (Acclaim PepMap 100, C18, 75 μm x 2 cm, 3 μm, 100 Å column) for 10 min (during which the column was held at 2% Solvent B). The peptides were resolved by switching the trap column to be in-line with the separating column, quickly increasing the gradient to 5% B over 5 min and then applying a 95 min linear gradient from 5% B to 25% B. Subsequently, the gradient was increased from 35% B to 40% B over 25 min and then increased again from 40% B to 90% B over 5 min. The column was then cleaned with a sawtooth gradient to purge residual peptides between runs in a sequence.

A Thermo Q-Exactive HF-X Orbitrap mass spectrometer was used to analyze protein digests. A full MS scan in positive ion mode was followed by 20 data-dependent MS scans. The full MS scan was collected using a resolution of 120000 (@ m/z 200), an AGC target of 3E6, a maximum injection time of 64 ms, and a scan range from 350 to 1500 m/z. The data-dependent scans were collected with a resolution of 15000 (@ m/z 200), an AGC target of 1E5, a minimum AGC target of 8E3, a maximum injection time of 55 ms, and an isolation window of 1.4 m/z units. To dissociate precursors prior to their reanalysis by MS2, peptides were subjected to an HCD of 28% normalized collision energies. Fragments with charges of 1, 6, 7, or higher and unassigned were excluded from analysis, and a dynamic exclusion window of 30.0 s was used for the data-dependent scans.

### Analyzing MS Data in FragPipe

FragPipe v20.0, along with MSFragger v3.8, IonQuant v1.9.8, and Philosopher v5.0, were used to analyze raw mass spectra with label-free quantification with match between runs enabled. Default settings were used except those delineated in Table 1 of the main text. Namely, the peptide digest pattern was set to semi-enzymatic, methionine oxidation and N-terminal acetylation were set as dynamic modification, and cysteine carbamidomethylation was set as a static modification. The workflows for *B. cereus* refolding and the reanalysis of PXD025926 were setup according to the conventions set forth in Figure S1B; wherein multiple LiP control-test pairs are normalized to a single TrP control-test pair. FragPipe outputs were the passed to FLiPPR with the command shown in Figure S1B. Output files from FLiPPR were then further processed in Python to create summaries and graphical representations of both data sets.

### Configuring FragPipe for LiP-MS Peptide Searches

**Figure S1.**
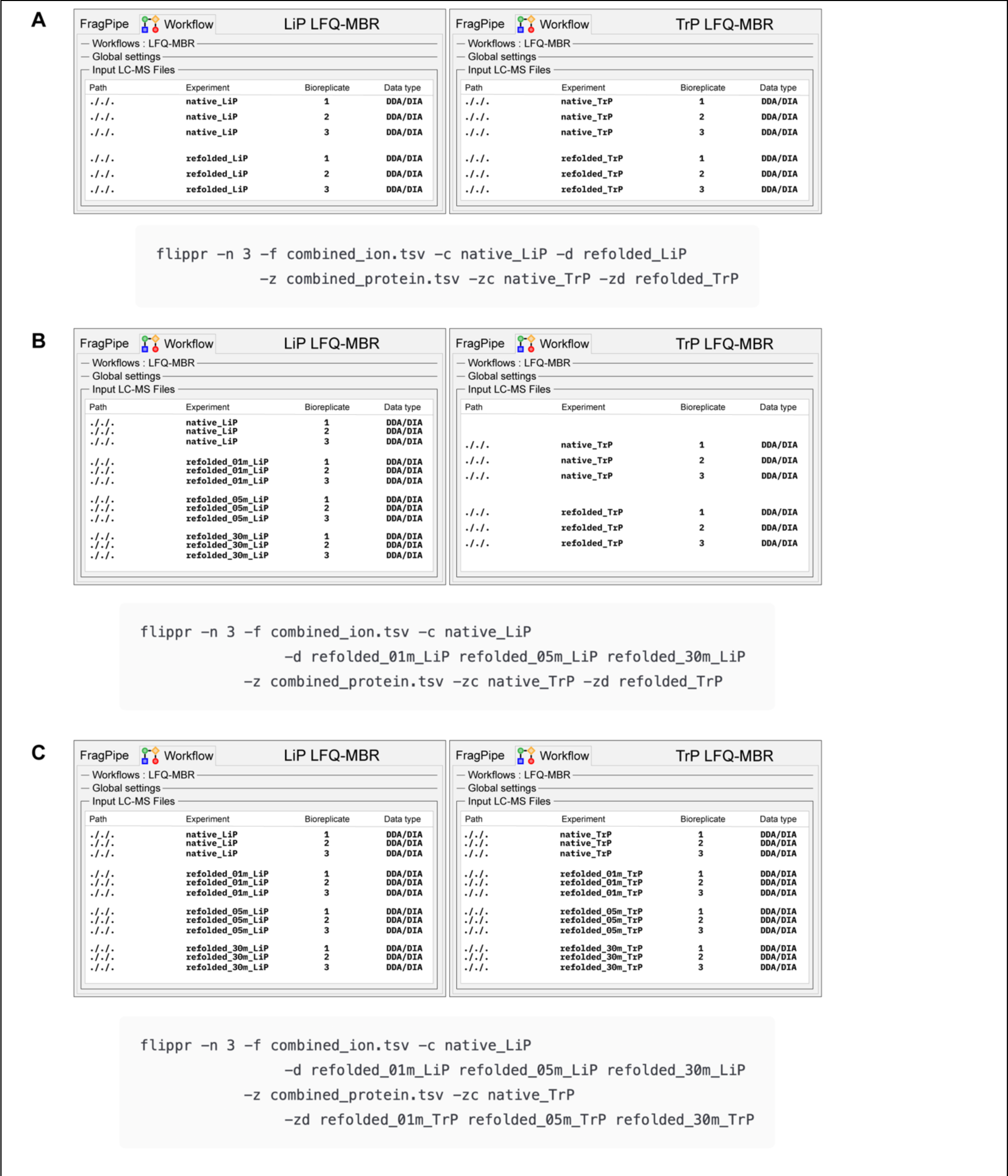
FragPipe LFQ-MBR workflows for LiP-MS experiments. Limited proteolysis experiments can be normalized in a large variety of ways. Here, three such scenarios are presented as standard conventions users are encouraged to use when analyzing LiP-MS data with FLiPPR. Each figure contains a schematic view of two separate FragPipe windows, each using the LFQ-MBR workflow, with the LiP experimental conventions shown on the left and TrP conventions shown on the right. Below these schematics, the FLiPPR command-line expression demonstrates how the LiP and TrP data must be passed into FLiPPR to achieve the expected analysis for the following scenarios: **(A)** Single LiP control-test pair normalized to a single TrP control-test pair. **(B)** Multiple LiP control-test pairs normalized to a single TrP control-test pair. **(C)** Multiple LiP control-test pairs normalized to an equal number of TrP control-test pairs.

